# ProFit-1D for quantifying J-difference edited data at 3T

**DOI:** 10.1101/2024.10.07.616795

**Authors:** Kimberly L. Chan, Tamas Borbath, Sydney Sherlock, Elizabeth A. Maher, Toral R. Patel, Anke Henning

## Abstract

Reproducible and accurate fitting of the magnetic resonance spectrum is critical for estimating metabolite concentrations. We have previously developed a fitting software called ProFit-1D which was shown to fit 9.4T semi-LASER data from the human brain with high accuracy and precision. In this study, we adapted ProFit-1D to fit J-difference edited spectra acquired at a clinical field strength of 3T and to assessed its performance in simulated and in vivo data. ProFit-1D was adapted to fit J-difference edited data with alterations to the fitting range to exclude the 1.3 ppm lipid resonance, starting T_2_ relaxation constants, initial fit parameters, and adaptive spectral baseline determination. The accuracy of ProFit-1D was systematically evaluated on simulated GABA-edited and 2-hydroxyglutarate-edited (2HG-edited) data with different types of in vivo parameter variations and compared to that of LCModel and Gannet, two software commonly used to fit J-difference edited data. The precision of ProFit-1D was also evaluated in GABA-edited spectra acquired in vivo in the occipital cortex (OCC) and medial prefrontal cortex (mPFC) of healthy participants at 3T using subsets of averages and compared to that of LCModel and Gannet. The 2HG fit error was also evaluated for ProFit-1D in 2HG-edited spectra acquired in glioma patients and compared to that of LCModel. Overall, it was found that ProFit-1D generally produced fits with low parameter fit errors across a variety of parameter variations. GABA, glutamate plus glutamine (Glx), and 2HG levels were also more accurately estimated with ProFit-1D than with LCModel and Gannet across different spectral disturbances and simulated concentrations. ProFit-1D was found to be as precise as LCModel and more precise than Gannet in estimating GABA and Glx. 2HG fit errors were 45% lower with ProFit-1D than with LCModel. Thus, ProFit-1D was found to produce high-quality fits to J-difference edited data with high accuracy and precision.

## Introduction

Proton magnetic resonance spectroscopy (^1^H-MRS) allows for the noninvasive quantification of neurochemicals in the brain which can provide biologically-relevant diagnostic and prognostic biomarkers as well as insight into the biological mechanisms underlying a variety of neurological and psychiatric conditions such as Parkinson’s disease(1,2), major depressive disorder(3,4), and brain tumors(5). ^1^H-MRS, however, suffers from a lack of spectral dispersion which results in significant overlap between different metabolite resonances. As such, there has been much research interest in the development of spectral fitting software that can accurately and reproducibly quantify individual metabolites. A variety of fitting methods have been developed to address this issue including AMARES(6), QUEST(7), Tarquin(8), and AQSES(9) which performs fitting in the time-domain, LCModel(10,11), Gannet(12), ProFit-2D(13,14), and Osprey(15), which performs fitting in the frequency-domain. Hybrid time-domain and frequency-domain approaches(16,17) and most recently, machine learning fitting algorithms(18,19), have also been previously proposed. Of these, LCModel is the most commonly-used spectral fitting tool and is considered the “gold standard” within the MRS community.

We have previously developed a fitting algorithm called ProFit-1D which uses adaptive spectral baseline determination and a novel cost function based on the frequency-domain residual, the time-domain residual, and the weighted spectral residual in the error minimization process(20). ProFit-1D was also shown to produce fits with slightly higher accuracy than that of LCModel and also high precision, which was however, slightly lower than that of LCModel. This version of the software, however, had the limitation of being optimized for and validated on high signal-to-noise ratio (SNR) sLASER data acquired on a 9.4T human scanner, of which only three such systems exist across the world. The software was also tested on data acquired in the occipital lobe of healthy subjects, which is an easy region to measure from due, in part, to its distance from air-tissue interfaces which would otherwise result in reduced shimming quality and consequently, broader linewidths, reduced SNR, and poorer water suppression.

Although sLASER and other non-edited MRS approaches are often sufficient to measure high-concentration metabolites with strong signals such as glutamate and N-acetylaspartate (NAA), the substantial spectral overlap hampers quantification of low concentration metabolites. As such, specialized techniques are often needed to simplify the MR spectrum so that signals from other compounds that may be overlapping a metabolite-of-interest are removed. This is often achieved through J-difference editing, which is considered the gold standard technique for detecting gamma-aminobutyric acid (GABA)(21,22), the major inhibitory neurotransmitter in the brain, and has been often used to quantify other low-concentration molecules such as N-acetylaspartate glutamate (NAAG)(23,24), glutathione (GSH) (25,26), lactate(26,27), and 2-hydroxyglutarate (2HG)(5), a MRS-detectable oncometabolite present in 80% of low grade gliomas(28) with important implications for diagnostic and prognostic purposes. In its initial release, ProFit-1D was optimized and demonstrated for non-edited MRS data, and does not have the capability to fit J-difference edited data as it relies on the presence of the large total choline (tCho), total creatine (tCr), and total NAA singlets in the spectrum to perform an initial fit and these singlets are often subtracted out in the difference spectrum. This first iteration of the error minimization process is also a necessary step in estimating the zero-order phase, first-order phase, and global frequency shift. Thus, the inability of ProFit-1D to fit J-difference edited data along with its initial application to MRS data acquired at the very uncommon field strength of 9.4T limits the use of this fitting software in its current form.

As such, the goal of this study was to adapt ProFit-1D to fit J-difference edited spectra acquired at a clinical field strength of 3T by making modifications to the fitting range, initial fit parameters, adaptive spectral baseline determination and assess its performance in simulated and in vivo data. Since the ground truth concentrations of in vivo spectra are unknown, the accuracy of the fitting software was systematically investigated in simulated GABA-edited and 2HG-edited spectra with known concentrations and data quality mimicking spectra acquired in vivo in healthy participants and brain tumor patients. Spectra with several possible perturbations including zero- and first- order phase changes, global frequency shifts, variable SNR, different spectral baselines, and variable amounts of Gaussian and Lorentzian line-broadening were simulated and the fit accuracy of ProFit-1D was evaluated and compared to that of LCModel(10) and Gannet(12), two commonly-used edited-MRS fitting software. The precision of ProFit-1D was then evaluated in GABA-edited data acquired in the occipital lobe (OCC) and medial prefrontal cortex (mPFC) of healthy participants and compared to that of LCModel and Gannet. The error in fitting the 2HG peak was also evaluated in 2HG-edited data acquired in glioma patients and compared between ProFit-1D and LCModel. No such evaluation was included for Gannet as the software only allows for the fitting of a few edited metabolites and does not include 2HG among its preset selections.

## Methods

### Edited ProFit-1D

Since J-difference editing is a subtraction method, the tCr and tCho singlets present in the conventional spectrum at 3.0 and 3.2 ppm, respectively, are missing from the difference spectrum. In the edited GABA and 2HG spectra, however, the editing pulse applied at 1.9 ppm results in suppression of the 2.0 ppm NAA peak in the edit-ON spectrum but not the edit-OFF spectrum and thus a fully-inverted NAA singlet is visible in the difference spectrum at 2.0 ppm. Thus, the first fitting iteration to estimate the zero- and first- order phases and global frequency shifts was limited to only the NAA peak. Additionally, the spectral fitting range was limited to 4.05-1.75 ppm to avoid complication from the lipid signals which can contaminate the spectra in the 1.75-0 ppm range(29). The fitted metabolites were also limited to only those present in the edited spectrum and the starting T2 relaxation constants were changed from 9.4T values to previously published 3T values (see simulated data for more information). The underlying spline baseline, which results from experimental imperfections such incomplete water suppression, has also been shown to significantly affect spectral quantification(30,31). In the first implementation of ProFit-1D, the spline smoothness was estimated using a previously published method(32) with parameters optimized for 9.4T data. As such, the maximum candidate baseline flexibility was changed from 10 to 7 effective dimensions per ppm while Akaike’s information criterion, which determines the smoothness factor, was changed from 15 to 5. These values were shown to be optimal for fitting 3T data by providing a good compromise between bias and variance (32).

The fitting software is freely available at https://gitlab.tuebingen.mpg.de/AG_Henning/ProFit-1D.

### Simulated data

Spatial simulations were performed in FID-A(33) across a 19 x 19 matrix for a (3.0 cm)^3^ voxel spanning 3.6 x 3.6 cm^2^ in the two refocusing pulse directions for GABA, 2-hydroxyglutarate (2HG), glutathione (GSH), glutamate (Glu), glutamine (Gln), N-acetyl-aspartyl-glutamate (NAAG), and N-acetylaspartate (NAA) with the same parameters as the in vivo data as well as Philips-specific refocusing pulse shape and duration, sequence timings, and phase cycling. The basis sets for ProFit-1D and LCModel fitting for the J-difference edited data included all the previously mentioned metabolites for fitting the 2HG-edited data but excluded 2HG for fitting the GABA-edited data. To best match in vivo conditions, GABA-edited spectra were simulated with concentrations extracted from LCModel fits to the OCC edit-OFF spectra with the exception of GABA which was taken from Gannet fits to the OCC difference spectra. 2HG-edited spectra were also simulated with concentrations extracted from LCModel fits to the edit-OFF spectra acquired in brain tumors with the exception of 2HG and GABA which were taken from ProFit fits to the tumor spectra. Both the GABA-edited and 2HG-edited data were simulated with a default 7 Hz Lorentzian linebroadening and base SNR of 310 for the GABA-edited data and 150 for the 2HG-edited data. The SNR was calculated as the absolute value of the total NAA peak (NAAG + NAA) divided by the root-mean-square of the noise. The SNR was lowered for the 2HG-edited spectra to simulate the lower SNR environment present in tumors (Supplementary Table 1).

To test the accuracy and precision of ProFit-1D, Gannet 3.3, and LCModel, edited GABA and 2HG spectra of in vivo data quality were simulated while varying only one parameter from equation 1 from (20) and keeping all others constant. These simulations included 15 different zero-order phases ranging from −35° to 35°, 15 different global frequency shifts ranging from −6 to 6 Hz, 17 different amounts of Gaussian line-broadening ranging from 2 to 18 Hz. The simulations also included different amounts of Lorentzian line-broadening (calculated as 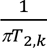 where T_2,k_ is the T_2_ relaxation constant for metabolite k) ranging from −2 to 2 times the standard deviation of previously reported T_2,k_ values(34–36). Since the T2 relaxation constant of 2HG is unknown, the initial T2 of 2HG was set to be the same as GABA (88 ms). Spectra were also simulated with 12 different spectral baselines which were created from cubic spline fits with manual definition of baseline points to previously-acquired GABA-edited spectra(22,37) and 15 different SNR values which ranged from ∼65 to ∼500 for the GABA-edited spectra and ∼35 to ∼160 for the 2HG-edited spectra. Spectral simulations were also performed with each metabolite concentration varied between −4 to 4 times the standard deviation plus the default values across all participants while holding the concentrations of all the other metabolites constant. Fifteen simulations were done for each metabolite to evaluate how accurately a large range of metabolite concentrations could be quantified. The simulated GABA-edited and 2HG-edited spectra with variations in the parameters are shown in Figure 1a and 1b, respectively. While the GABA-edited spectra was fit with all three software, the 2HG-edited spectra was only fit with ProFit-1D and LCModel as Gannet cannot fit 2HG-edited spectra. For LCModel, the spectra were fit from 1.95 to 4.2 ppm with SPTYPE = ‘mega-press-3’, and did not include any a priori knowledge regarding metabolite ratios.

**Figure 1.**
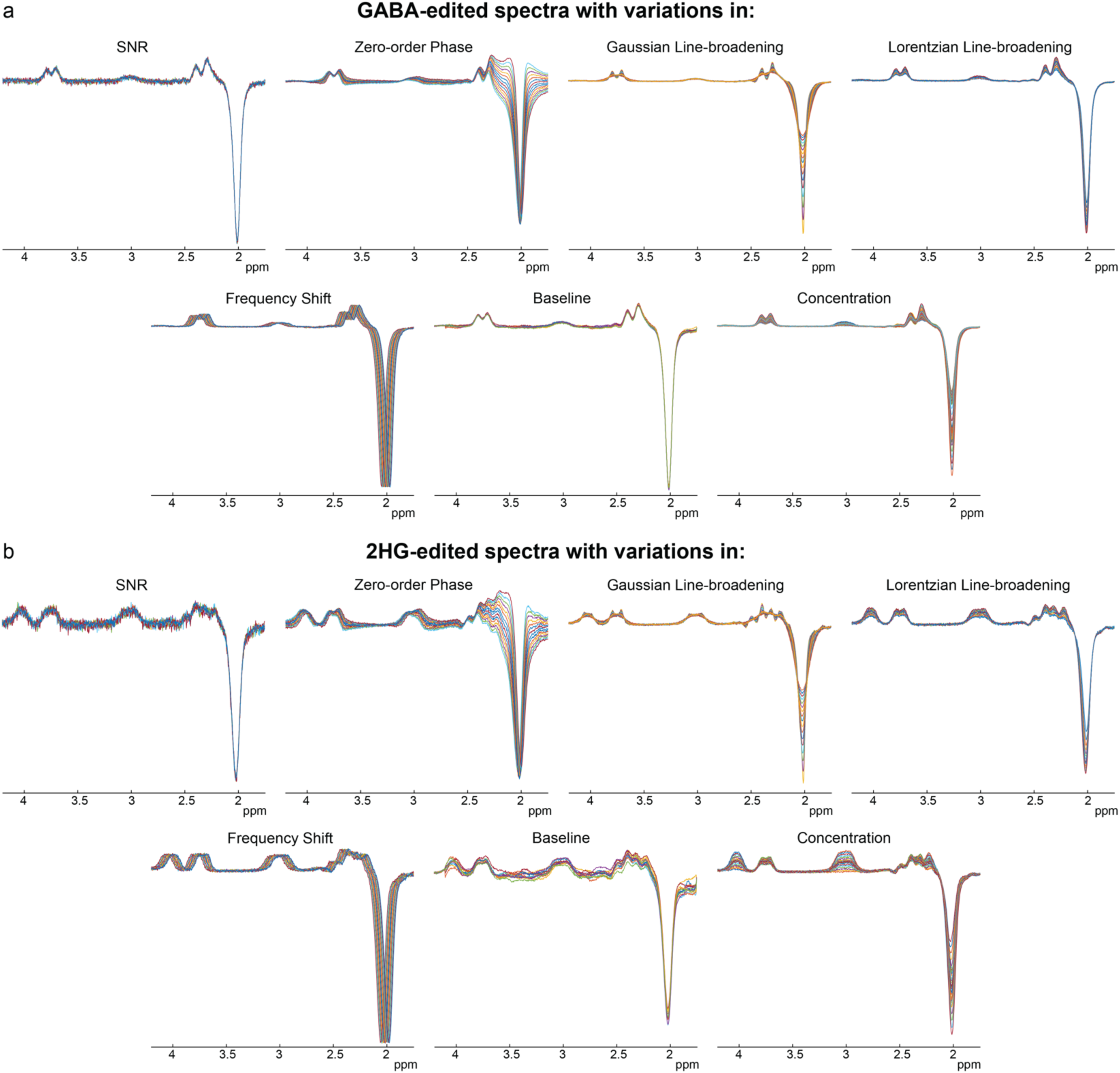
Simulated GABA-edited spectra (a) and 2HG-edited spectra (b) with different parameter variations that were used to evaluate ProFit-1D, Gannet, and LCModel. Twelve to fifteen simulated spectra were generated per parameter variation setup.

### In vivo data

In vivo data were acquired on a Philips Achieva 3T after participants gave informed written consent with local Institutional Review Board approval. MRS data were acquired with a MEGA-PRESS sequence, repetition time (TR) of 2 s, 2048 spectral points, and spectral bandwidth of 2 kHz. GABA-edited data were acquired in 12 healthy volunteers (6 female, 6 male, age 24.8 ± 2.3 years) in a (3 cm)^3^ region in the occipital lobe (OCC) and in 8 healthy volunteers (6 female, 2 male, age 26 ± 2.6 years) in a 2.5 x 3.5 x 3 cm^3^ region in the medial prefrontal cortex (mPFC). Other scan parameters included 320 transients, an echo time (TE) of 80 ms, an editing pulse duration of 20 ms, an edit-ON pulse at 1.9 ppm, and an edit-OFF pulse at 1.5 ppm. 2HG-edited data were acquired in 5 glioma patients (5 female, age 41.6 ± 8.1 years) with a TE of 70 ms, TR of 2 s, an editing pulse duration of 15 ms, an edit-ON pulse at 1.9 ppm, an edit-OFF pulse at 7.5 ppm, and

352 transients. All edited-MRS scans were performed with prospective frequency correction during the scan based on the frequency of a non-suppressed water reference scan collected once every 20 averages as described previously(38). In the glioma patients, a T2-weighted FLAIR was also acquired for voxel localization. The voxel varied in location and was either (3.5 cm)^3^ or (3 cm)^3^ depending on the tumor size. Spectral SNR and linewidths for the in vivo GABA-edited spectra and the 2HG-edited spectra are shown in Supplementary Table 1.

Data were preprocessed using Gannet 3.3(12) with retrospective phase and frequency correction performed in the time domain(39). To evaluate the precision in fitting the in vivo GABA-edited spectra, two test-retest spectra (160 averages each) were created from all available spectral averages per subject, fit with Gannet 3.3, LCModel, and ProFit-1D, and used to calculate coefficients-of-variation (CoV). The error in fitting 2HG was also evaluated for both ProFit-1D and LCModel and calculated as the root-mean-square of the fit residual in the 4.2-3.8 ppm range divided by the maximum of the 2HG peak at 4.02 ppm. No test-retest analysis was performed for the 2HG-edited spectra due to the inherently-low SNR of the data which makes it difficult to measure accurate CoVs from spectra reconstructed from subsets of all averages.

## Results

The accuracy of the fitted ProFit-1D parameters resulting from systematic parameter changes are shown in Figures 2a-e for the GABA-edited spectra and Figures 2f-j for the 2HG-edited spectra. Overall, Lorentzian line-broadening, Gaussian line-broadening, and SNR influenced the fit accuracy of the GABA-edited and 2HG-edited spectra the most. However, the fits to the GABA-edited spectra generally displayed high accuracy with parameter fit errors of < |10°| for the zero-order phase, < |5°| for the first-order phase, < |0.5| Hz for the Gaussian line-broadening parameter, and < |1| Hz for the global frequency shift. Relative to the GABA-edited spectra, the ProFit-1D fits to the 2HG-edited spectra are less accurate, however, the fits still generally displayed high accuracy with parameter fit errors of < |6°| for the first-order phase and < |0.5| Hz for the Gaussian line-broadening parameter.

**Figure 2.**
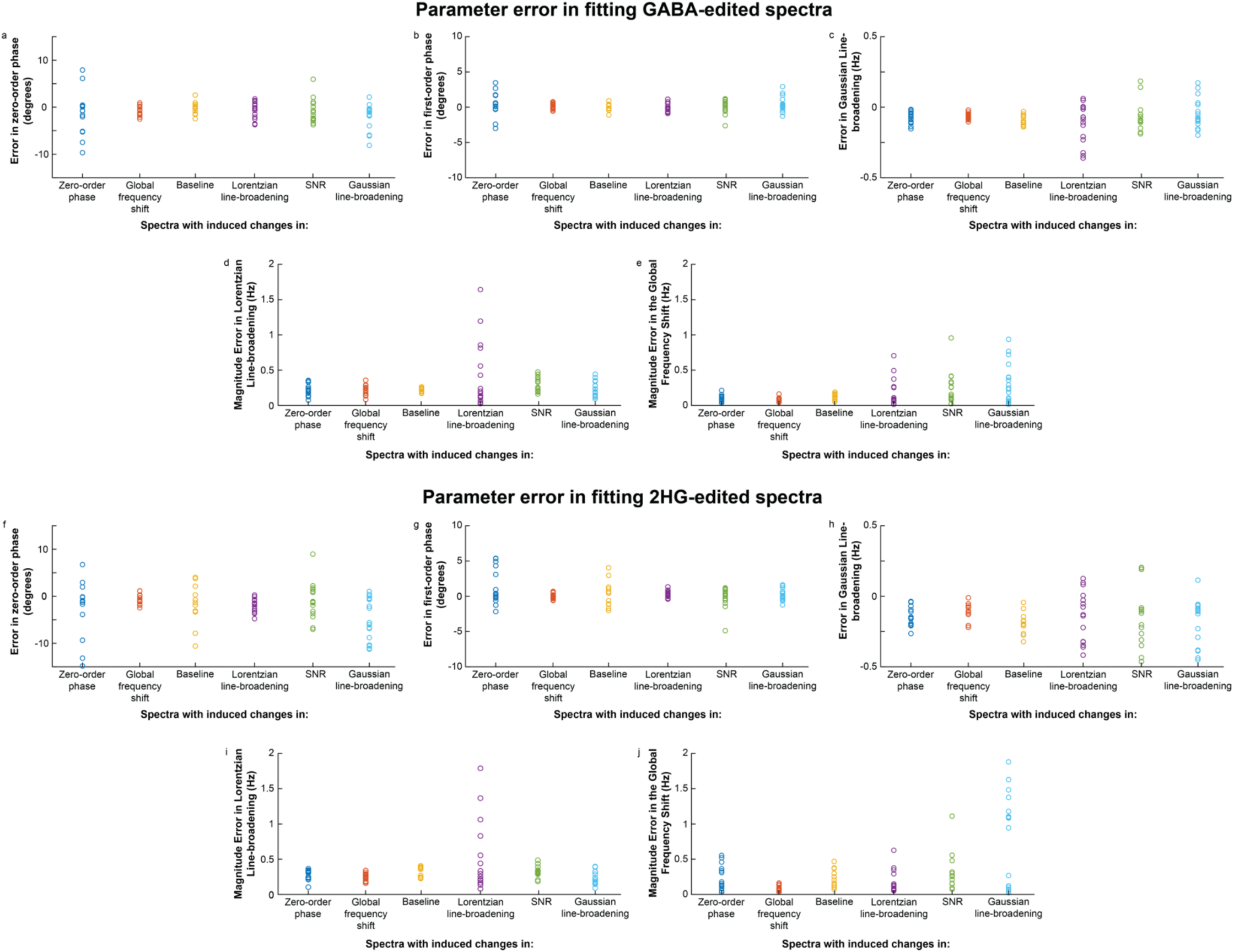
Evaluation of ProFit-1D parameter estimation error in zero-order and first-order phase, Gaussian and Lorentzian linebroadening, and global frequency shift in GABA-edited spectra (a-e) and 2HG-edited spectra (f-j) with different parameter variations (Figure 1). Overall, fits to both the GABA- edited and 2HG-edited spectra displayed high accuracy with generally low parameter fit errors for all spectral simulation setups.

Percent changes in fitted metabolite concentrations for each simulated parameter variation are shown in Figures 3a-e for the GABA-edited spectra. Generally, ProFit-1D measures GABA and Glx more accurately from the GABA-edited spectra than LCModel and GABA more accurately than Gannet. While the GABA and Glx concentrations estimated with ProFit-1D were fairly accurate across all parameter variations, changes in Gaussian line-broadening appears to influence the accuracy of ProFit-1D metabolite concentrations the most with a minor effect on the Glx concentrations and large effect on NAA and NAAG concentrations. Of all the metabolites fitted with ProFit-1D, GSH was the least accurately estimated across all spectral disturbances. In addition, GABA concentrations, as estimated by all three fitting software, appear to be most sensitive to baseline distortions, zero-order phase changes, and SNR variation. However, the GABA concentrations estimated with Gannet appear to be the most sensitive to these three parameter variations. Although the Glx concentrations estimated with LCModel are fairly accurate across all parameter variations, the GABA concentrations estimated with LCModel are somewhat inaccurate across all parameter variations. In general, all 3 fitting software performed best with the frequency shift variation versus other parameter variations.

**Figure 3.**
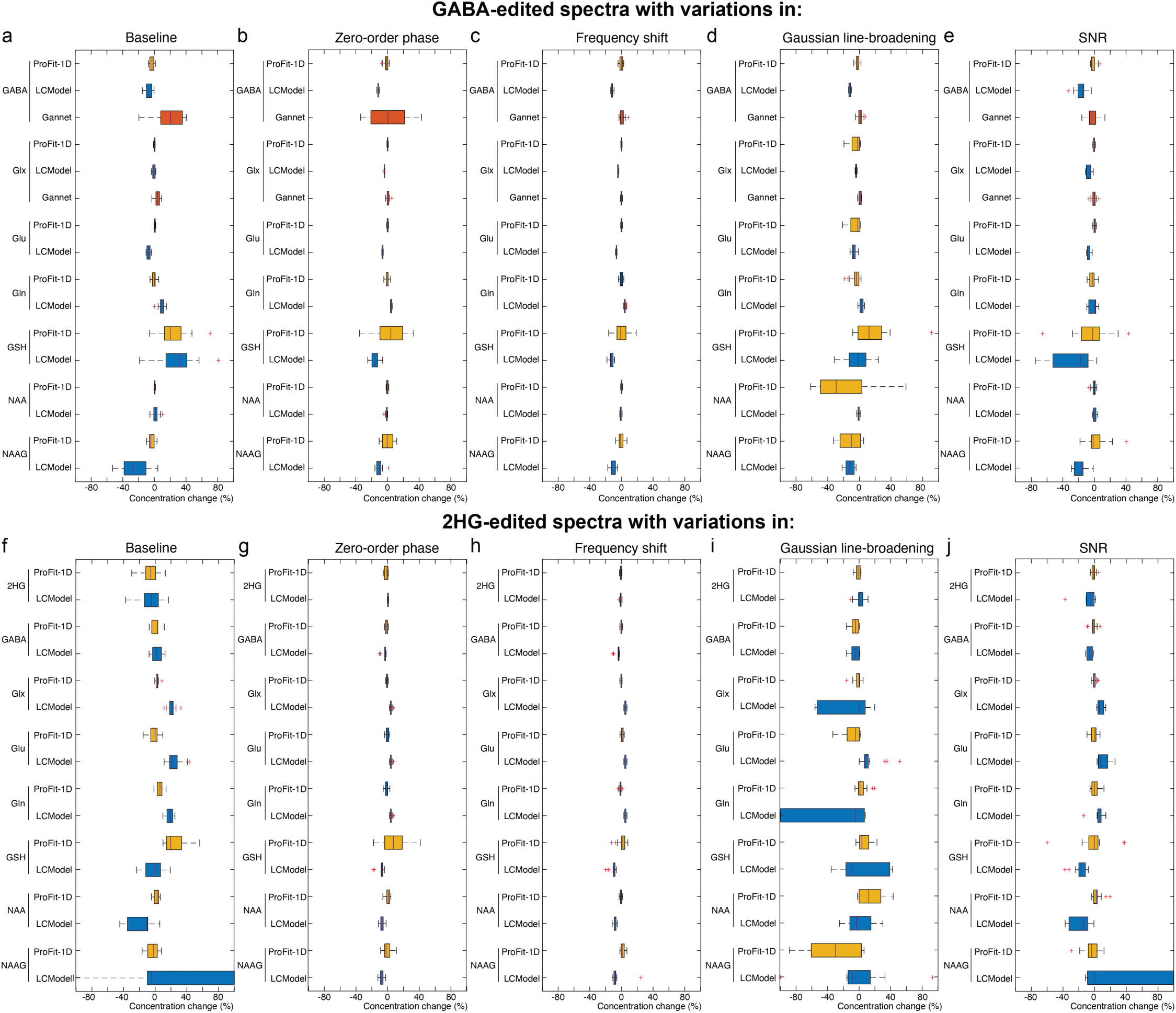
Percent change in fitted concentrations with different parameter changes are shown in (a-e) for the simulated GABA-edited spectra and in (f-j) for the simulated 2HG-edited spectra. Although Glx was estimated from the GABA-edited spectra relative well across all spectral perturbations for all 3 fitting software, GABA was estimated from the GABA-edited spectra more accurately with ProFit-1D than with Gannet or LCModel. In the simulated 2HG-edited spectra, ProFit-1D estimates of 2HG, GABA, and Glx are generally more accurate than that of LCModel. ProFit-1D estimates of the other metabolites, NAA, NAAG, Gln, and Glu, are also generally more accurate than that of LCModel for both the GABA-edited spectra and 2HG-edited spectra with the exception of GSH.

Percent changes in fitted metabolite concentrations for each simulated parameter variation are shown in Figures 3f-j for the 2HG-edited spectra. Overall, ProFit-1D is more accurate than LCModel when measuring 2HG, GABA, and Glx concentrations across all parameter variations. As in the GABA-edited spectra, metabolite concentrations fitted using ProFit-1D and LCModel are least sensitive to frequency shifts. In addition, metabolite concentrations fitted using LCModel, are most sensitive to changes in Gaussian line-broadening with estimated Glx concentrations that changed substantially (up to 52%) with this parameter change. Metabolite concentrations fitted using ProFit-1D, are also most sensitive to changes in Gaussian line-broadening, but less so in general than those estimated with LCModel.

Correlation plots between the fitted metabolite concentrations versus simulated concentrations from the GABA-edited spectra are shown in figure 4a. For the GABA-edited spectra, ProFit-1D was just as accurate as Gannet in estimating Glx and GABA concentrations and slightly more accurate than LCModel in estimating GABA, Glx, NAAG, and GSH concentrations. Figure 4b shows the correlation plots between the fitted metabolite concentrations and simulated concentrations estimated from the 2HG-edited spectra. Overall, ProFit-1D was found to be substantially more accurate than LCModel in estimating metabolite concentrations from the 2HG-edited spectra. In particular, LCModel was found to moderately overestimate Glx and Gln levels at all simulated concentrations, misestimate Glu at all concentrations, especially at lower concentrations (< 1.25 mM), misestimate NAAG and GSH at all concentrations, and underestimate NAA at higher concentrations (2 mM). Although the performance of LCModel was improved for 2HG and GABA, LCModel was found to underestimate 2HG at lower concentrations (< 2.5 mM) and underestimate GABA at higher concentrations (> 2.5 mM).

**Figure 4.**
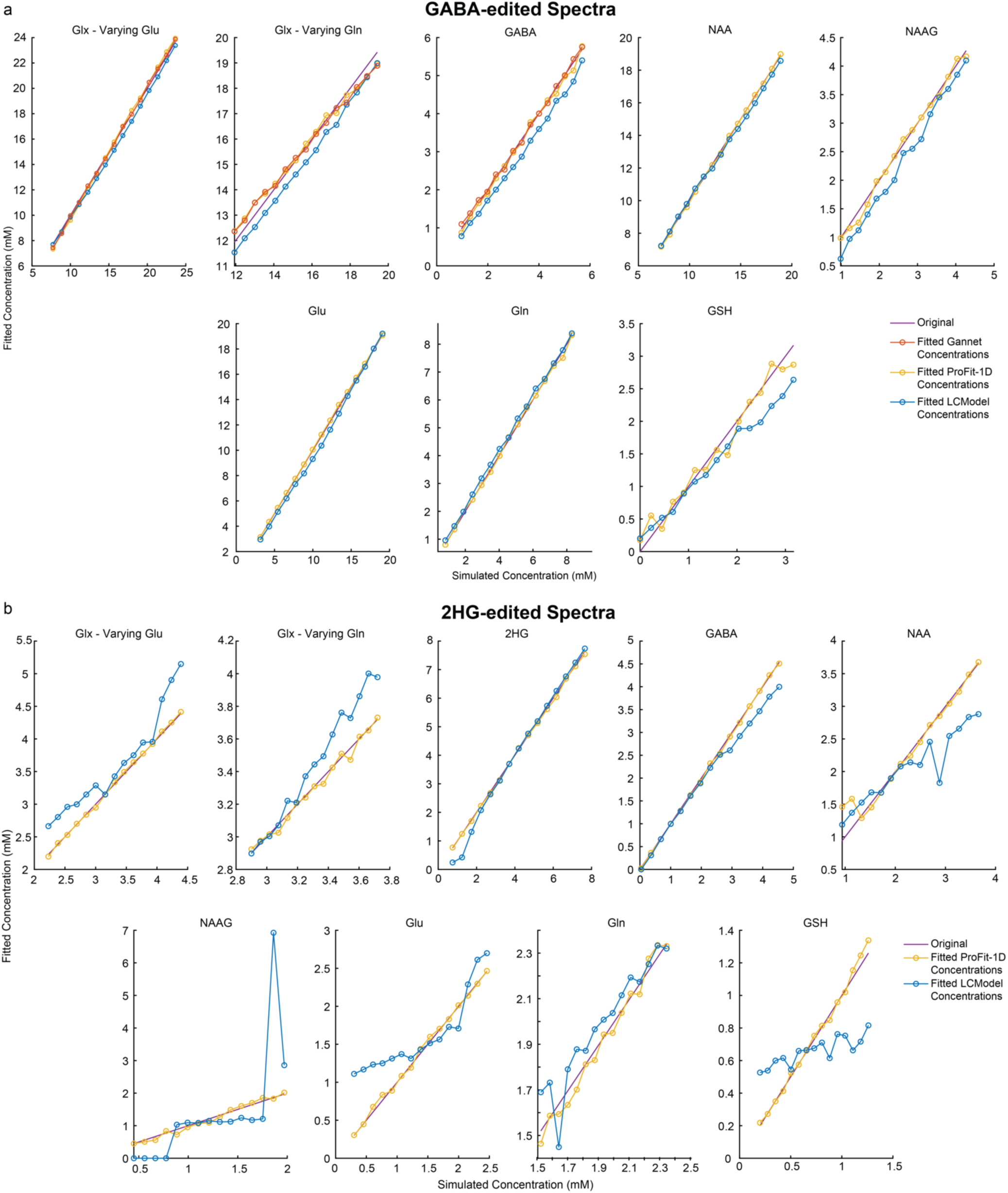
Fitted concentrations versus simulated concentrations for all measured metabolites in the GABA-edited (a) and 2HG-edited (b) spectra. In the GABA-edited spectra, ProFit-1D and Gannet fit GABA and Glx with similarly high accuracy and both ProFit-1D and Gannet fit GABA and Glx slightly more accurately than LCModel. ProFit-1D was also found to fit NAAG, Glu, Gln, and GSH in the GABA- edited spectra slightly more accurately than LCModel. In the 2HG-edited spectra, ProFit-1D was found to fit Glx, 2HG at low values, and GABA at high values more accurately than ProFit-1D. ProFit-1D also generally fits the other metabolites (NAA, NAAG, Glu, Gln, and GSH) more accurately than LCModel.

Figure 5 shows LCModel, Gannet, and ProFit-1D fits to the in vivo GABA-edited MRS data acquired in the OCC and mPFC of healthy participants. Although the spectra are noticeably noisier in the mPFC (Figure 5b) than in the OCC (Figure 5a), all three fitting software demonstrated high-quality fits to the spectra in both regions. It can be seen, however, that Gannet does not fit other co-edited metabolites such as NAA and NAAG and the Glx peak fitted by Gannet contains a component from GSH in addition to Glu and Gln. Figure 5c shows the in vivo CoVs for GABA and Glx estimated the GABA-edited spectra acquired from the OCC of healthy participants. Although all three fitting software were found to have high precision, it can be seen that the GABA and Glx concentrations as measured with LCModel and ProFit are more precise than those measured with Gannet. In addition, LCModel fits GABA most precisely with a low mean CoV of 5.1% and ProFit fits Glx most precisely with a low mean CoV of 3.8%. Figure 5d shows the in vivo CoVs for GABA and Glx estimated from GABA-edited spectra acquired in the mPFC of healthy participants where it can be seen that the precision in fitting GABA and Glx in the mPFC is lower than that in the OCC for all three fitting software. Similar to what was found in the OCC, however, Gannet is less precise than LCModel and ProFit-1D in fitting GABA with a mean CoV of 24%. ProFit-1D was found to fit GABA most precisely with a mean CoV of 14.4% and LCModel was found to fit Glx most precisely with a mean CoV of 8%. Altogether, ProFit and LCModel were found to be equally precise in fitting the main metabolites, GABA and Glx, with comparable average CoVs of 8.5% and 8.6% for ProFit and LCModel, respectively, across both regions. On the other hand, the average CoVs for GABA and Glx across regions were 12% for Gannet which was 40% higher than that for ProFit and LCModel.

**Figure 5.**
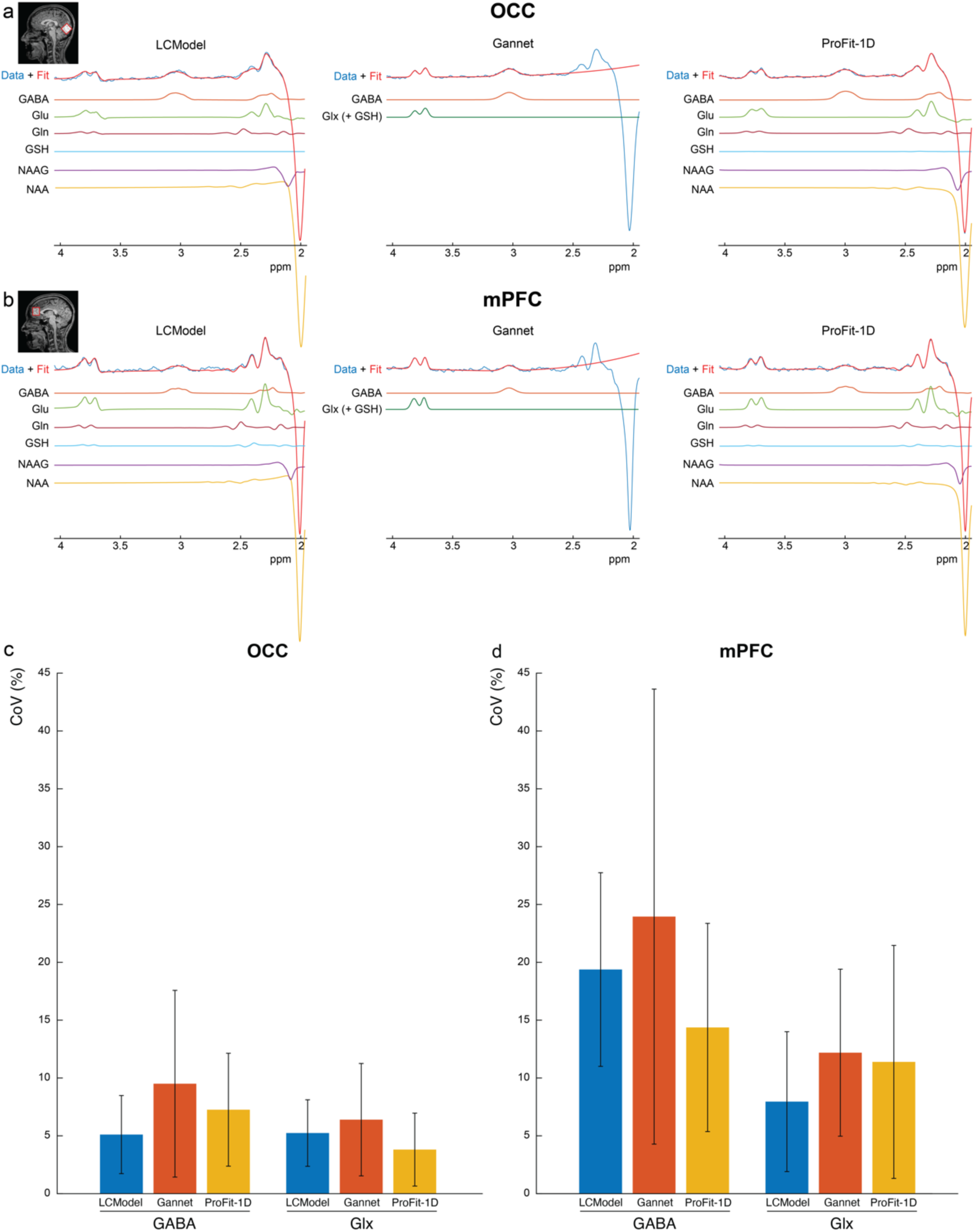
In vivo GABA-editing results. Representative ProFit-1D, Gannet, and LCModel fits to the in vivo GABA-edited spectra showing high-quality fits for all fitting software in are shown in (a) for the OCC and (b) for the mPFC. Glu, Gln, GSH, NAAG, and NAA fits were only performed for LCModel and ProFit as Gannet does not include separate fits for these metabolites. Instead, the 3.0 ppm GABA peak is modeled as a single Gaussian and the 3.75 ppm peak is modeled as two Gaussians. As such, Gannet’s ‘Glx’ peak also contains a small contribution from GSH. Bar plots of in vivo GABA and Glx CoVs from fits to the GABA-edited spectra are shown for the OCC (c) and mPFC (d). CoVs were higher in the mPFC than in the OCC. Relative to Gannet, both LCModel and ProFit-1D had lower CoVs for GABA and Glx in both regions while LCModel and ProFit-1D displayed relatively equal CoVs across both regions for GABA and Glx.

Supplementary figure 1 shows the CoVs for the fitted concentrations for the other metabolites, NAAG, NAA, GSH, Glu, and Gln, in the GABA-edited spectra. As with the main metabolites, GABA and Glx, the CoVs were generally higher for all but Gln. In addition, ProFit-1D and LCModel were found to be equally precise with average CoVs of 16% for ProFit-1D and 18% for LCModel over the other metabolites. This is true for the individual metabolites as well with average ProFit-1D CoVs of 9.4% for NAAG, 2.2% for NAA, 18% for GSH, 8.8% for Glu, and 43% for Gln and average LCModel CoVs of 11% for NAAG, 2.5% for NAA, 18% of GSH, 7.9% for Glu, and 39% for Gln.

The results from the ProFit-1D and LCModel fits to the in vivo 2HG-edited spectra are shown in Figure 6. High-quality fits to the in vivo 2HG-edited spectra can be seen for both LCModel and ProFit-1D in Figure 6a, however, the overall fit appears to be more accurate with ProFit-1D than with LCModel. This is corroborated by the results in Figure 6b where it can be seen that the average 2HG fit error is 45% higher with LCModel than with ProFit-1D.

**Figure 6.**
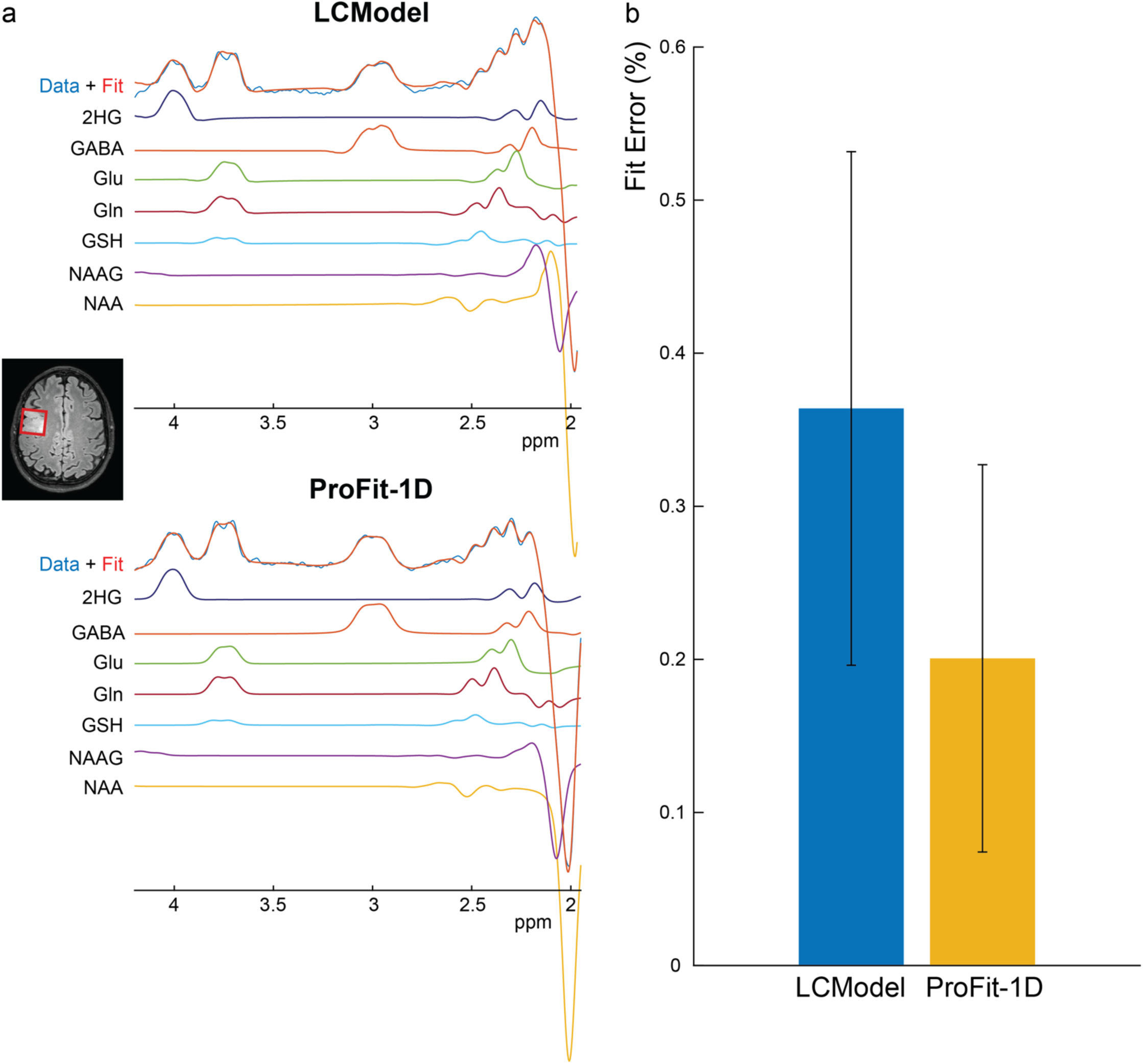
In vivo 2HG-editing results. Example voxel placement overlayed on T2-weighted FLAIR image and representative ProFit-1D and LCModel fits to the 2HG-edited spectra showing high-quality fits are shown in (a). Gannet was not used to fit the 2HG-edited spectra as it does not have the capability of doing so. A bar chart showing significantly higher 2HG fit error with LCModel than with ProFit-1D is shown in (b).

## Discussion

In this study, ProFit-1D was adapted to fit J-difference edited data acquired at a clinical field strength of 3T and evaluated for accuracy and precision in simulated data and in vivo 2HG- and GABA- edited data acquired in gliomas and healthy participants, respectively, and compared to that of Gannet and LCModel in the GABA-edited data and that of LCModel in the 2HG-edited data. Overall, ProFit-1D was found to be more accurate and precise than Gannet and more accurate than LCModel in fitting the GABA-edited and 2HG-edited spectra. ProFit-1D was also found to be more precise than Gannet and as precise as LCModel for both the main metabolites, GABA and Glx, as well as the other metabolites, NAA, NAAG, Glu, Gln, and GSH.

Here, the accuracy of the fitting software was performed on simulated data sets as the ground truth metabolite concentrations and spectral properties of in vivo data are unknown. As a part of this test, a systematic evaluation of disturbances of the edited ^1^H-MRS, such as phase distortion, frequency shifts, baseline distortions, line broadening and noise, which mimicked in vivo conditions was performed to identify possible areas of fitting underperformance. In general, similar to what was shown previously in the original implementation, ProFit-1D did not have any systematic errors in estimating fitting parameters or metabolite concentrations across all possible spectral disturbances. In addition, estimation of the main metabolites, GABA, Glx, and 2HG, remained accurate across all disturbances. ProFit-1D had the highest uncertainty when fitting GSH across most of the parameter variations. This is likely due to the inherently low concentration of GSH combined with the low editing efficiency of GSH (when performing GABA-editing) and high spectral overlap with other metabolites, particularly Glu and Gln. In addition, ProFit-1D had the highest uncertainty when fitting the highly-overlapped metabolites in edited spectra with different amounts of Gaussian linebroadening. This was particularly true for GSH as well as NAA and NAAG which are heavily overlap by each other in the 2HG- and GABA-edited spectrum. Regardless, ProFit-1D had the highest overall accuracy among the three fitting software across all spectral perturbations, especially when considering the main metabolites of interest, GABA, Glx, and 2HG.

ProFit-1D was also found to estimate changes in GABA and Glx concentration just as accurately as Gannet and more accurately than LCModel when using the default spectral parameters. Overall, the differences in accuracy between LCModel and ProFit-1D from fits to the GABA-edited spectra were small and the inaccuracy of LCModel was consistent across simulated concentrations. In the 2HG-edited spectra, however, the fitted metabolite concentrations using LCModel were significantly more inaccurate and not consistent across all concentrations, most notably, when estimating 2HG at lower concentrations. This discrepancy in fit performance between GABA-edited and 2HG-edited spectra is likely due to the 50% lower SNR of the 2HG-edited spectra relative to the GABA-edited spectra as the LCModel fits were found to be more sensitive to changes in SNR. This non-uniform misestimation of metabolite levels with LCModel could limit the ability to accurately diagnose IDH-mutant gliomas with low 2HG concentrations, and hamper the ability to monitor metabolite levels in response to treatment.

In addition to these accuracy tests in simulated data, the precision of ProFit-1D was evaluated on in vivo GABA-edited data sets acquired in both the OCC and mPFC and compared to that of Gannet and LCModel. Overall, Gannet was the least precise of the three fitting software when fitting GABA and Glx and ProFit-1D and LCModel were found to be equally precise for all metabolites. This is in contrast to the original ProFit-1D implementation in sLASER acquired at 9.4T which showed slightly lower precision than LCModel for the harder-to-fit metabolites. It should be noted that in Gannet’s the 3.0 ppm and 3.75 ppm peak in the GABA-edited spectrum are directly fit and assigned to ‘GABA’ and ‘Glx’, respectively. As such, it should be noted that Gannet’s ‘Glx’ measurement also contains a moderate contribution from GSH which takes up almost a fifth of 3.75 ppm peak with a median (interquartile range) of 18% (4.1%) as estimated from ProFit-1D fits to the in vivo GABA-edited spectra. This introduces a confound to the ‘Glx’ concentrations as measured with Gannet which may be especially amplified in situations where GSH is elevated such as in post-traumatic stress disorder(40), mild cognitive impairment(41), depression(42). As such, this extra contamination from co-edited GSH in addition to Glx in the 3.75 ppm peak is a drawback of using lineshape models to fit the edited spectra versus a linear combination of simulated spectral patterns from contributing metabolites. Additionally, with this fitting approach, the other GABA resonance at 2.28 ppm and Glx resonances at 2.11, 2.33 and 2.43 ppm are not included in the fit. Thus, Gannet’s fitting approach may provide more flexibility than LCModel and ProFit-1D, both of which use all of the GABA and Glx resonances in the fitting algorithm, which may have resulted in the reduced precision relative to the two other fitting software.

To date, most validation and optimization of spectral fitting pipelines have been performed on data acquired in healthy controls(12,15,20) and in regions with favorable measurement conditions(43) (e.g. high SNR, low linewidths). Here, ProFit-1D was evaluated in a variety of different regions and volunteer populations. Optimization and validation of spectral fitting software in more challenging regions such as the mPFC and in patient populations, such as brain tumor patients, is important for clinical applicability. The mPFC is a more difficult region to measure from due to its proximity to susceptibility effects originating from the nearby air-tissue interface which can result in larger linewidths, poorer water suppression, and reduced SNR. Consistent with this, the metabolite concentrations measured from the mPFC generally had higher CoVs, and hence, lower precision from all three fitting software. Relative to scanning this region in healthy volunteers, scanning brain tumors in patients is even more challenging due to the inherently lower SNR environment and even more challenging shimming conditions from the adjacent metal implants put in place after surgical resection, which can cause significant baseline distortions. Despite these challenges, ProFit-1D was able to produce high-quality fits to both the 2HG-edited and GABA-edited data.

It should be noted that this revision of ProFit-1D was tested on macromolecule-suppressed (MM-suppressed) edited GABA data, but not edited GABA data with co-edited macromolecules (GABA+). To date, the co-edited MM peak underlying GABA in the GABA+ signal is poorly characterized and not much is known about its composition, coupling constants, and multiplet pattern(44). While some prior work have modeled the MM component of 3.0 ppm GABA+ peak as a single Gaussian(43), there is evidence that this MM component is mostly composed of lysine, whose multiplet at 3.0 ppm is more asymmetric in nature(45). In addition, lysine has another resonance at ∼1.3 - 1.7 ppm which would not be entirely saturated by the editing pulse at 1.9 ppm, assuming a typical pulse duration of 14 ms and bandwidth of 88 Hz, and which could help improve fitting of the edited spectrum if properly modeled. As such, future work will focus on exploring the properties of this underlying lysine/MM peak(45), examining best ways of incorporating its model into ProFit’s fitting algorithm, possibly, by using an experimentally-measured MM-baseline which has been shown to improve non-edited spectral fits(30,31), and evaluating the impact this modeling has on the ProFit-1D GABA measurements.

Additionally, ProFit-1D was optimized in this study for GABA-edited and 2HG-edited spectra. While the tCr and tCho singlets are subtracted out in the GABA- and 2HG-edited spectrum, a fully-inverted NAA singlet at 2.0 ppm remains. This singlet is needed in the first iteration of ProFit-1D to estimate the zero-order phase, first-order phase, and global frequency shift. Despite the absence of the two other singlets, tCr and tCho, ProFit-1D was able to fit the GABA-edited and 2HG-edited data well. This is not always the case with other edited MRS acquisitions, however. Although J-difference editing is most commonly applied to detect GABA, other low-concentration metabolites such as NAAG(23,24), GSH(25,29,46), and lactate(26,27) have also been commonly detected with J-difference editing. In the edited spectra for these metabolites, all three singlets (tCr, tCho, and tNAA) are subtracted out. Thus, to fit these spectra with ProFit-1D, alternate strategies are needed for the first iteration fit to estimate initial global parameters and avoid overfitting and local minimal solutions. Future work will focus on implementing this for the difference spectra of other editable metabolites by possibly estimating these global parameters from the edit-OFF spectra or the sum (edit-ON + edit-OFF) spectra.

## Conclusions

In this study, a newly-developed spectral fitting software, ProFit-1D, which was originally adapted for sLASER data acquired at 9.4T, was modified to fit J-difference edited data acquired at a clinical field strength of 3T. The accuracy and precision of ProFit-1D was then evaluated in simulated and in vivo GABA- edited and 2HG-edited data. The GABA-edited data was acquired in the OCC and mPFC of healthy volunteers while the 2HG-edited data were acquired from brain tumors of glioma patients. ProFit-1D was found to be more accurate and precise than Gannet and more accurate than LCModel in fitting the GABA- edited spectra and the 2HG-edited spectra. Additionally, ProFit-1D was found to be just as precise as LCModel in fitting all metabolites.

## Acknowledgements

This work was funded by the Cancer Prevention and Research Institute of Texas (CPRIT) (Grant Number: RR180056).

## Notes

**Grant sponsors**: This project was sponsored by the Cancer Prevention and Research Institute of Texas (CPRIT) / Grant number: RR180056.

https://gitlab.tuebingen.mpg.de/AG_Henning/ProFit-1D

### Competing Interest Statement

The authors have declared no competing interest.

