## supplementary information for "ProFit-1D for quantifying J-difference edited data at 3T"

| | GABA-edited spectra (healthy patients) | | 2HG-edited spectra (glioma patients) |
|---|---|---|---|
| | OCC | mPFC | |
| SNR | 226 (71) | 165 (44) | 120 (110) |
| NAA Linewidths (Hz) | 6.8 (1) | 5.4 (1.5) | 9.8 (12.1) |

**Supplementary Table 1.** MRS data quality metrics including the SNR, FWHM of the 2.0 ppm NAA peak. Values are expressed as median (interquartile range). GABA-edited spectra acquired from healthy normal participants had higher SNR and lower spectral linewidths than the 2HG-edited data acquired in glioma patients.

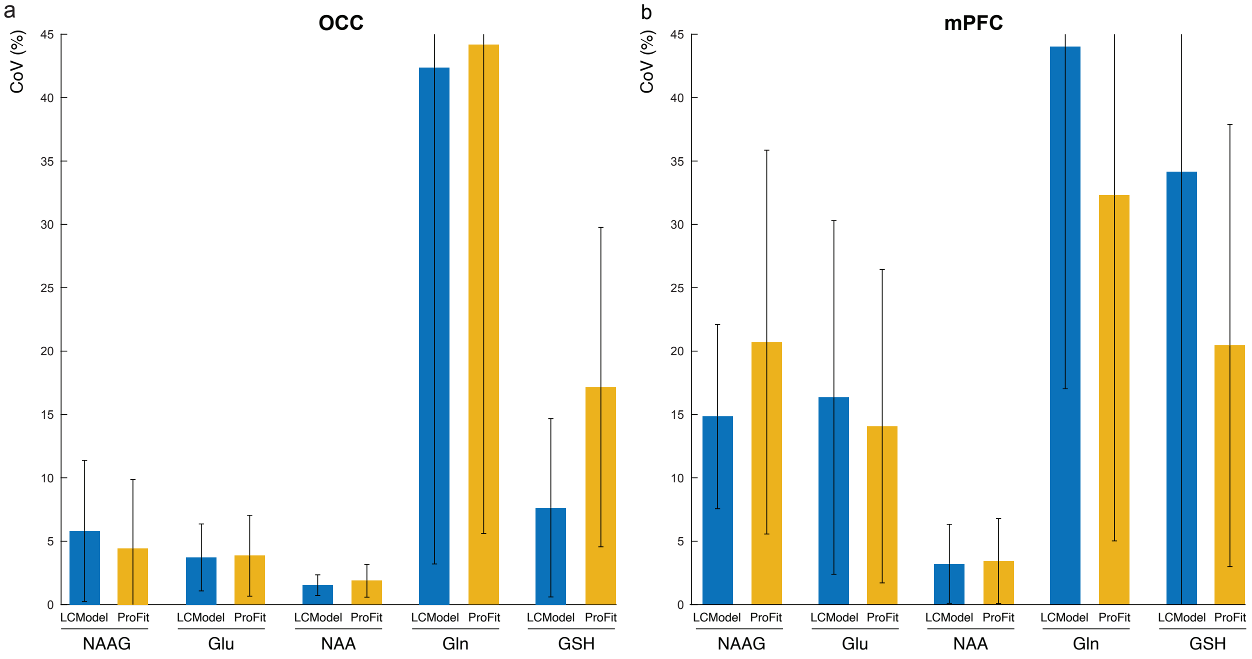


**Supplementary Figure 1.** In vivo CoVs of NAAG, Glu, NAA, Gln, and GSH concentrations estimated from ProFit-1D and LCModel fits to the GABA-edited spectra. Over both regions, ProFit-1D and LCModel were equally precise with comparable CoVs which ranged from ~2% for NAA to 45% for Gln.
